# Biobank-scale inference of multi-individual identity by descent and gene conversion

**DOI:** 10.1101/2023.11.03.565574

**Authors:** Sharon R. Browning, Brian L. Browning

**Author notes:** Address for correspondence: SRB; BLB.

## Abstract

We present a method for efficiently identifying clusters of identical-by-descent haplotypes in biobank-scale sequence data. Our multi-individual approach enables much more efficient collection and storage of identity by descent (IBD) information than approaches that detect and store pairwise IBD segments. Our method’s computation time, memory requirements, and output size scale linearly with the number of individuals in the dataset. We also present a method for using multi-individual IBD to detect alleles changed by gene conversion. Application of our methods to the autosomal sequence data for 125,361 White British individuals in the UK Biobank detects more than 9 million converted alleles. This is 2900 times more alleles changed by gene conversion than were detected in a previous analysis of familial data. We estimate that more than 250,000 sequenced probands and a much larger number of additional genomes from multi-generational family members would be required to find a similar number of alleles changed by gene conversion using a family-based approach.

## Introduction

Segments of identity by descent (IBD) are shared tracts of DNA that have been inherited from a recent common ancestor who has lived within the past few hundred generations. These segments can be detected in genotype data from population samples by looking for long tracts of identical-by-state alleles that are present in two or more haplotypes.^1^

Detected IBD segments are useful in many applications,^2; 3^ including IBD mapping,^4-7^ detecting signatures of natural selection,^8; 9^ identifying close relatives,^10-12^ and estimating kinship coefficients,^13^ demographic history,^14^ effective population size,^15; 16^ migration rates,^17^ mutation rates,^18; 19^ and recombination rates.^20; 21^

IBD segment detection is often applied to pairs of haplotypes or pairs of individuals.^1; 2; 5; 9; 12; 22-28^ However, the pairwise approach has limitations. One major limitation is the quadratic scaling of the number of pairwise IBD segments with sample size. The number of pairwise IBD segments in large data sets is enormous. For example, a recent analysis of UK Biobank SNP array data with 408,883 individuals found 77.7 billion segments of IBD with length > 2 cM.^9^ In contrast, the time required to detect and the space required to store multi-individual IBD using our proposed method scales linearly with sample size. A second limitation of the pairwise approach is that it is difficult to make use of additional information that comes from considering multi-individual IBD in downstream analyses. Multi-individual IBD can help distinguish genotype error from recent mutation and gene conversion^19; 29^ and it can help distinguish between certain familial relationships.^30^

Instead of pairs of haplotypes, we work with clusters of haplotypes. A set of haplotypes form an IBD cluster at a locus if they have a recent common ancestor. Moving along the chromosome, haplotypes may leave or join the IBD cluster at points of crossovers. Whereas a pairwise IBD segment is defined by two haplotypes along with the beginning and end positions of the IBD segment, multi-IBD needs a different approach. For small data sets, one can record the set of haplotypes in each IBD cluster and the genomic positions at which the cluster membership changes.^6^ However, this approach is unwieldy for large data sets. We use an alternate approach. At each position of interest, we record the cluster containing each haplotype. Thus, the IBD cluster data is like phased genotype data, but with IBD clusters replacing alleles. Indeed, our software outputs the IBD cluster information in a format that is intentionally similar to Variant Call Format (VCF)^31^ for genotype data (Figure S1), with haplotype cluster indices replacing allele indices. Since it is not necessary to output IBD clusters at each genotyped variant, the output file storing the IBD clusters can have many fewer lines than the input VCF file containing the genotype data. The number of columns in the output file increases linearly with sample size, in contrast to the number of pairwise IBD segments which can increase quadratically with sample size.

An important property of multi-individual IBD that is typically ignored when analyzing pairwise IBD is transitivity: if haplotypes *h*_1_ and *h*_2_ share a recent common ancestor X, and haplotypes *h*_2_ and *h*_3_ share a recent common ancestor Y, then haplotypes *h*_1_ and *h*_3_ must share a recent common ancestor who will be X or Y (Figure 1A). However, defining pairwise IBD in terms of a minimum shared segment length results in non-transitivity because the additional pairwise IBD segments implied by transitivity may be smaller than the length threshold (Figure 1B). In order to obtain transitivity of IBD, we must accept that real pairwise IBD segments imputed by transitivity may be relatively short in some cases. An additional problem arising when inferring IBD is that it is difficult to determine the precise endpoints of IBD sharing.^9^

**Figure 1:**
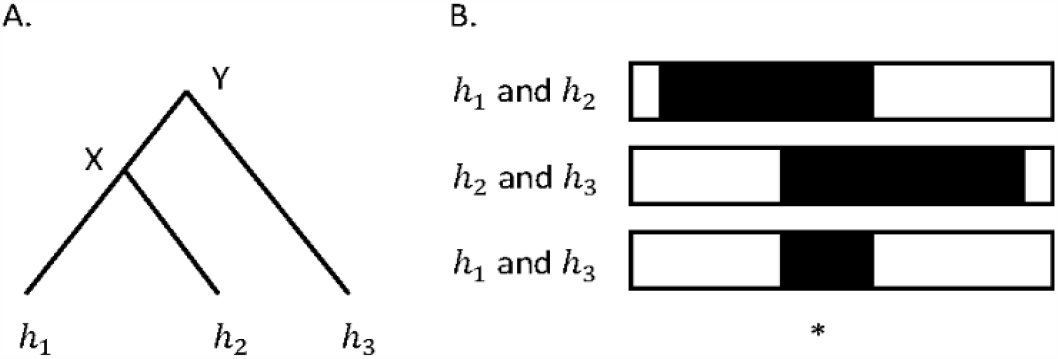
Transitivity of IBD. **A**. The coalescent tree relationship at a given point in the genome is shown for haplotypes *h*_1_, *h*_2_, and *h*_3_ which are mutually IBD at this location. Haplotypes *h*_1_ and *h*_2_ have common ancestor X, while haplotypes *h*_1_ and *h*_3_, and haplotypes *h*_2_ and *h*_3_, have common ancestor Y. **B**. Pairwise IBD status (black for IBD, white for non-IBD) is shown for the three pairs of haplotypes along a region of the chromosome around the focal position (denoted *). The IBD extends to either side of the focal point until reaching a point of recombination on one of the ancestral lineages. Although the IBD sharing between haplotypes *h*_1_ and *h*_2_, and between haplotypes *h*_2_ and *h*_3_, is long and may exceed a pre-defined length threshold, the IBD between haplotypes *h*_1_ and *h*_3_ is relatively short and may not meet the length threshold for pairwise IBD sharing.

Estimating segment endpoints that go beyond the actual extent of IBD and then imputing additional IBD to obtain IBD transitivity can lead to incorrect clustering (Figure 2A). Previous approaches to determining multi-individual IBD have generally looked for highly-connected clusters of pairwise IBD, and then added or removed some pairwise IBD to obtain IBD transitivity.^6; 32; 33^ This approach scales quadratically with cluster size. In this work, we solve the problem by trimming a fixed genetic distance (e.g. 1 cM), from each end of the pairwise identity by state (IBS) segments before imputing additional IBD to obtain IBD transitivity (Figure 2B). Although our approach is equivalent to trimming pairwise IBD segments, our use of IBD transitivity results in computation time scaling linearly, rather than quadratically, with sample size.

**Figure 2:**
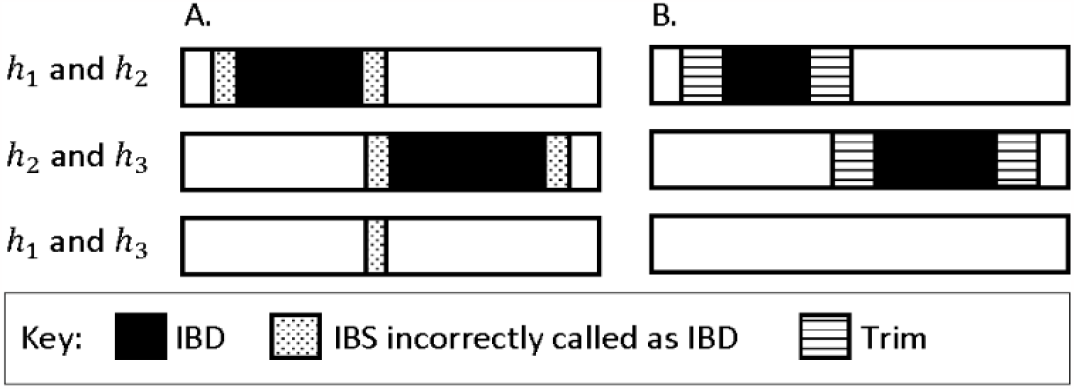
IBD transitivity with and without trimming IBS segments. IBD and IBS in a genomic region is shown for the three pairings of three haplotypes (haplotypes *h*_1_, *h*_2_, and *h*_3_). **A**. The IBD between haplotypes *h*_1_ and *h*_2_ is derived from a different recent common ancestor than that for the IBD between haplotypes *h*_2_ and *h*_3_. IBS that is not due to the recent common ancestors is incorrectly called as IBD at the ends of the IBD segments. As a result, transitivity leads to a region of IBS being incorrectly called as IBD between haplotypes *h*_1_ and *h*_3_. **B**. A trim is applied to the ends of the pairwise IBS regions and no IBD is called between haplotypes *h*_1_ and *h*_3_.

In previous work we showed that multi-individual IBD is useful for distinguishing genotype error from recent mutation or gene conversion.^19; 29^ Homologous gene conversion occurs during meiosis on the transmitted haplotype when the haplotype that is copied from the parent is interrupted by a short tract copied from the parent’s other haplotype. When this occurs, the transmitted haplotype’s allele is changed at any position at which the parent is heterozygous in the copied region. These allele changes can create discordant alleles within an IBD cluster if the gene conversion arose since the most recent common ancestor.

Because gene conversion tracts are small, with length generally less than 1400 base pairs,^34^ gene conversion is difficult to study. One approach to studying gene conversion uses sperm typing.^35^ Another approach uses multi-generational family data. The use of multi-generational families (as compared to the use of nuclear families) helps to resolve phase and to distinguish true gene-conversion allele changes from genotype errors.^34; 36^ Two additional approaches do not directly detect gene conversion allele changes but estimate rates of gene conversion, using IBD or linkage disequilibrium.^18; 37^ Previous studies using these approaches have shown that gene conversion tracts in humans are on the order of hundreds of kilobases in length,^34; 35^ the genome-wide average rate at which sites are included in gene conversion tracts is approximately 6 × 10^−6^ per bp per meiosis,^18; 34; 36^ and gene conversions hotspots tend to coincide with crossover hotspots.^34-36^

In this work, we show how multi-individual IBD can be used to infer instances of alleles that have been changed by recent gene conversion. In contrast to existing methods, our method can be applied to population samples rather than family data or sperm-typing, and it can infer changed alleles rather than rates. Application of our method to UK Biobank sequence data detects more than 9 million allele conversions.

## Methods

### Multi-individual IBD

We say that two haplotypes *h*_1_ and *h*_2_ are *confidently* identical by descent at marker *m* if the two haplotypes share an identical allele sequence that is at least *L* cM in length and if the shared allele sequence contains marker *m* after trimming *T* cM from each end of the shared sequence. The default values in our ibd-cluster software program for *L* and *T* are *L* = 2 cM and *T* = 1 cM, which we found to give accurate results in our analyses of simulated data and UK Biobank sequence data (see Results).

We say two haplotypes *h* and 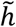 are identical by descent at marker *m* if there is a sequence of haplotypes, *h*_1_, *h*_2_, …, *h*_*n*_ such that 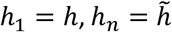, and for which each pair of consecutive haplotypes in the sequence is confidently identical by descent at marker *m*. This is an equivalence relation that defines a partition of the haplotypes at a marker. We call this partition the IBD partition at the marker. The sets in the IBD partition are clusters of haplotypes. Each set contains one or more haplotypes. Two haplotypes are in the same set of the partition at a marker if and only if the two haplotypes are identical by descent at the marker.

### Algorithm

We first exclude markers with low minor allele frequency (MAF). All IBD clustering analyses in this study exclude markers with MAF < 0.1. This retains the most informative markers and reduces the number of discordant alleles caused by genotype error and recent mutation in IBD segments. If a variant has multiple alleles, we define the MAF to be the second largest allele frequency.

We then aggregate sets of consecutive, closely spaced markers to form multi-allelic markers. The alleles of each aggregate marker are the allele sequences at the constituent markers. We construct aggregate markers by processing individual markers in chromosome order. We aggregate the first marker of an aggregate marker with any following markers that are within a user-specified distance (0.005 cM by default). Each marker is part of exactly one aggregate marker. The base coordinate of an aggregate marker is the median of the base coordinates of the constituent markers. The genetic position of the aggregate marker base position is obtained by linear interpolation from the input genetic map. We exclude markers that are outside the boundaries of the genetic map because extrapolation of genetic distances is inaccurate outside the genetic map. We perform IBD clustering at the aggregate marker positions. For convenience, we use the term marker to refer to an aggregate marker when describing the IBD clustering algorithm.

We assume that input phased genotypes contain *H* haplotypes with indices 1, 2, …, *H*, and *M* markers in chromosome order with indices 1, 2, …, *M*. We use a sliding marker window with length ≥ *L* cM. The first window begins with the first marker and ends with the first marker that is at least *L* cM away. The last marker advances by one marker per window. The first marker in a window is the closest marker that precedes the last marker by at least *L* cM.

Let *W*_1_, *W*_2_, …, *W*_*J*_ be the sequence of windows. For each window, *W*_*j*_, let 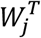 denote the trimmed window that is obtained by trimming *T* cM (*T* ≤ *L*/2) from each end of *W*_*j*_. We can restate our previous definition of confidently identical by descent in terms of marker windows: two haplotypes are confidently identical by descent at marker *m* if there is a window *W*_*j*_ in which the haplotypes have the same alleles and *m* is contained in the trimmed window 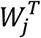. Our IBD clustering algorithm assigns all haplotypes that are confidently identical by descent at marker *m* to the same IBD cluster. At each marker, the clustering algorithm initially places each haplotype in a separate IBD cluster (one haplotype per cluster). The clustering algorithm merges two IBD clusters at a marker whenever there is a pair of haplotypes, one from each cluster, that are confidently identical by descent at the marker.

We use the Positional Burrows-Wheeler Transform (PBWT)^38^ to identify lists of haplotypes that share the same allele sequence in each window. The PBWT’s computation time scales linearly with the number of markers and with the number of haplotypes.^38^ If a list of haplotypes with the same allele sequences is identified in window *W*_*j*_, we merge the IBD clusters containing these haplotypes at each marker in the trimmed window 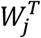 since the haplotypes are confidently identical by descent at the markers in the trimmed window. Due to IBD transitivity, it is only necessary to merge IBD clusters for adjacent pairs of haplotypes in the list. It is not necessary to merge IBD clusters for all pairs of haplotypes.

We use a disjoint-set data structure to perform computationally efficient merging of IBD clusters at a marker.^39^ The disjoint-set data structure stores a set as a tree in which each node is labeled with a unique element of the set. Two IBD clusters can be merged by setting the root node of one tree to be a child of the root node of the other tree. Additional efficiency is gained through the use of path compression and union by rank to reduce the depths of the trees.^39^

Our algorithm’s memory requirements scale linearly with the number of haplotypes, *H*. The use of a sliding marker window limits the number of disjoint set data structures that must be stored in memory. Technical details and pseudocode for the clustering algorithm are included in the Supplementary Methods.

### Detection of allele conversions

Allele conversions are instances in which an allele on a haplotype carried by one or more individuals in the data has been changed due to a gene conversion occurring in an ancestral meiosis. If two identical-by-descent haplotypes carry different alleles at a position, the difference could be due to an allele conversion, but it could also be due to genotype error or mutation.

We use multi-individual IBD to distinguish between genotype error and allele conversions. We infer an allele conversion in an IBD cluster if at least two haplotypes in the cluster share one allele and at least two haplotypes share a different allele. If the allele differences in this cluster were due to genotype error it would require two genotype errors at the same position in the same IBD cluster, which is a low probability event if genotype errors occur independently. Since we require that an IBD cluster must have at least two copies of each of two alleles in order to infer an allele conversion, an IBD cluster must contain at least four haplotypes in order for an allele conversion to be detected.

In order to reduce the problem of mutation creating apparent allele conversions, we consider only markers with MAF greater than 5% when detecting allele conversions. This MAF filter is applied to the input markers and is independent of the MAF filter used in the IBD clustering. It is not necessary to use the same markers for IBD detection and allele conversion detection because IBD clusters change slowly along the chromosome and can be extrapolated to nearby positions. We use the IBD clustering at the aggregate marker that is closest to the site at which we are detecting allele conversions. Non-recurrent recent mutations will have frequency much lower than 5% in a large sample that has relatively few closely-related individuals, and thus will be excluded when detecting allele conversions. Allele conversions at markers with MAF below 5% will not be detected, and this needs to be considered in downstream analyses. However, most allele conversions occur at markers with higher frequency because allele conversions can only occur when the ancestral individual in whom the gene conversion occurred was heterozygous.

IBD clusters carrying an uncalled deletion can lead to false-positive allele conversion inferences. Genotypes carrying uncalled deletions are typically miscalled as homozygote genotypes for the non-deleted allele. These miscalled deleted alleles may cause an IBD cluster to appear to contain more than one allele. To account for deletions miscalled as homozygotes when calling allele conversions, we ignore IBD clusters for which all individuals carrying haplotypes in the cluster are homozygous. This step will remove some real allele conversions due to all the individuals being homozygous by chance.

Gene conversion creates discordant alleles in IBD segments. If we include such positions in the multi-individual IBD detection, our algorithm won’t detect the IBD segments and hence won’t find the allele conversions. Consequently, we run multiple IBD clustering and gene conversion analyses that use different sets of markers and then combine the results. In each IBD clustering analysis, we include data across 9 kb, then leave a gap of 9 kb, so that there is a repeating pattern of length 18 kb. To ensure coverage of the whole genome, and to avoid edge effects at the ends of the gaps, we perform three analyses on each chromosome, with the 18 kb regions offset by 0, 6, and 12 kb (Figure S2). In each analysis, we first perform IBD clustering at the aggregate markers. After IBD clustering, we detect allele conversions at individual markers in the 9 kb gaps. When detecting allele conversions at a marker in one of the gaps, we use the IBD clustering for the nearest aggregate marker in terms of genetic distance. Within each 9 kb gene gap, we ignore 1.5 kb on each end when detecting allele conversions to ensure that a gene conversion tract in an analyzed region does not overlap with one of the surrounding 9 kb IBD detection regions. The length of a gene conversion tract is less than 1.5 kb with high probability.^35^ After removing these 1.5 kb end regions, 6 kb remains for each offset, and each point in the genome is included in one of the three analyzed 6 kb gene conversion detection regions.

### Accounting for end-of-chromosome effects

When reporting IBD rates, we consider only positions that are more than *L* − *T* cM from the ends of the chromosomes and simulated regions. Since we trim *T* cM from the end of each IBS segment, no IBD will be detected in the first and last *T* cM on a chromosome. There will be reduced IBD detection between *T* and *L* − *T* cM from an end of a chromosome because any trimmed IBS segment that is contained within the first or last *L* − *T* cM of a chromosome will correspond to an untrimmed IBS segment that is too short to meet the IBS length threshold.

### Simulated data

We simulated 20 regions of size 10 Mb, each with 10,000 simulated individuals, and we simulated an additional 20 regions of size 10 Mb, each with 125,000 simulated individuals. The smaller simulated data sets are used to measure the accuracy of IBD clustering and allele conversion detection because it is possible to extract the true IBD and allele conversions in these data (see below). The larger data sets have a similar number of individuals to the UK Biobank sequence data that we analyze. We used msprime v1.2 to perform the simulation.^40^ We used a constant recombination rate of 1 cM/Mb, a mutation rate of 1.5 × 10^−8^ per basepair per meiosis, and gene conversion with an initiation rate of 0.02 per Mb and mean tract length of 300 bp. We simulated an exponentially growing population with initial size of 10,000 and growth of rate 3% per generation for the past 200 generations, for a final size of 3.7 million.

We added uncalled deletions to the simulated data. It is assumed that uncalled deletions will tend to have low frequency in real data. We thus convert 1% of the simulated variants with MAF less than 1% into uncalled deletions. Each deletion extends from the converted variant in the direction of increasing position for an exponentially distributed distance with mean 300 bp. Within this range, alleles on the deletion haplotype are set to be equal to the alleles on the individuals’ other haplotype, thus making the individuals homozygous within the deletion. Although each deletion has frequency less than 1%, the genotypes changed to homozygous genotypes may be from SNPs having any MAF.

We added genotype error at a rate of 0.02%. Each genotype had a 0.02% probability of having a randomly chosen allele changed. This error rate is based on the discordance rate seen in TOPMed sequence data.^9; 41^ We excluded variants with MAF <= 0.01, removed the phase information, and used Beagle 5.4 to statistically phase the genotypes.^42^

### Determination of false discovery rates in simulated data

We recorded the simulated ancestry trees for the smaller simulations (10,000 individuals) and used the ancestry trees to determine the true IBD and allele conversions as described in Supplementary Methods.

IBD false discovery rates were determined as follows: At each position in the output IBD cluster data, we determined the identical-by-descent haplotype pairs that are implied by the clustering (i.e., all pairs of haplotypes within the same IBD cluster at that position). We then checked the true IBD status for each of the inferred identical-by-descent haplotype pairs at that position. We summed the numbers of false positives identical-by-descent haplotype pairs across all positions and divided by the number of interrogated haplotype pairs, summed across all positions.

Inferred allele conversions involve an allele that has been inferred to have been changed in two or more haplotypes in an IBD cluster at the locus due to gene conversion. For each inferred allele conversion, we determined whether the corresponding IBD cluster contains a pair of haplotypes with different alleles which is included in the list of true allele conversions. To obtain the allele conversion false discovery rate, we summed the number of false positives across all inferred allele conversions and divided by the number of inferred allele conversions.

### UK Biobank data

We analyzed whole autosome sequence data from 125,361 individuals from the UK Biobank.^43^ These are the White British individuals from the initial release of 150,119 sequenced genomes.^44^ The 150,119 genomes were phased using Beagle 5.4.^42; 45^ The deCODE genetic map was used for IBD clustering.^46^

## Results

### Simulated data

In the smaller simulated data (10,000 individuals) we assessed accuracy when varying the minimum pairwise IBS length, *L*, from 1.5 to 3 cM and varying the IBS segment trim, *T*, from 0.1 to 1.0 cM (see Table S1). We did not consider larger values of *L* and *T* because these data sets are not large enough to find much IBD or many allele conversions for larger values for these parameters. We find that IBD and allele conversion false discovery rates are low for most of the settings considered. As expected, false discovery rates and detection rates decrease when *L* or *T* is increased.

We analyzed the larger simulated data that contains 125,000 individuals with *L* = 2 and *T* = 1, because these parameters give relatively high rates of IBD and allele conversion detection in the smaller simulated data while having a relatively low IBD false discovery rate and allele conversion false discovery rate (8.0 × 10^−5^ and 0.010 respectively; Table S1). The average pairwise IBD rate in the larger data was 6.6 × 10^−6^ per haplotype pair per locus which is similar to that in the smaller data (6.0 × 10^−6^; Table S1). A total of 284,838 allele conversions were detected across the 200 Mb of simulated larger data.

Analysis of subsets of the 125,000 individuals demonstrates that the software scales linearly in computation time and memory requirements with sample size (Figure S3). Analysis of 120,000 sequenced individuals required only 6 GB of memory.

### UK Biobank data

We analyzed the autosomal UK Biobank White British data with minimum IBD length threshold *L* = 2 cM and trim *T* = 1 cM. Inferring the IBD clusters for chromosome 1 on a compute node with 96 cores (DNAnexus instance type: mem2_ssd1_v2_x96) took 205 minutes for a single gap offset. Most of this computation time is spent reading, parsing, and filtering the input VCF records. The chromosome 1 data has more than 31 million variants, of which only 151 thousand remain after filtering to remove markers with MAF<0.1 and markers in the gap regions. If we analyze an input file containing only the variants with MAF>0.05, the compute time for marker filtering and inferring IBD clusters is reduced to only 11 minutes.

The average pairwise IBD rate was 3.4 × 10^−5^ per haplotype pair per locus. On average, 41.4% of haplotypes were members of a cluster of size 1 at a marker, and 37.8% of haplotypes were members of a cluster of size 4 or larger (Figure 3). On average there are 10,878 clusters per marker that had at least 4 haplotypes. We detected 9,313,066 allele conversions across the autosomes. The number of inferred allele conversions at a MAF is proportional to the heterozygosity as expected for a relatively homogeneous population (Figure S4), which gives confidence that a high proportion of the inferred allele conversions result from actual gene conversions rather than recurrent mutations or other artifacts.

**Figure 3.**
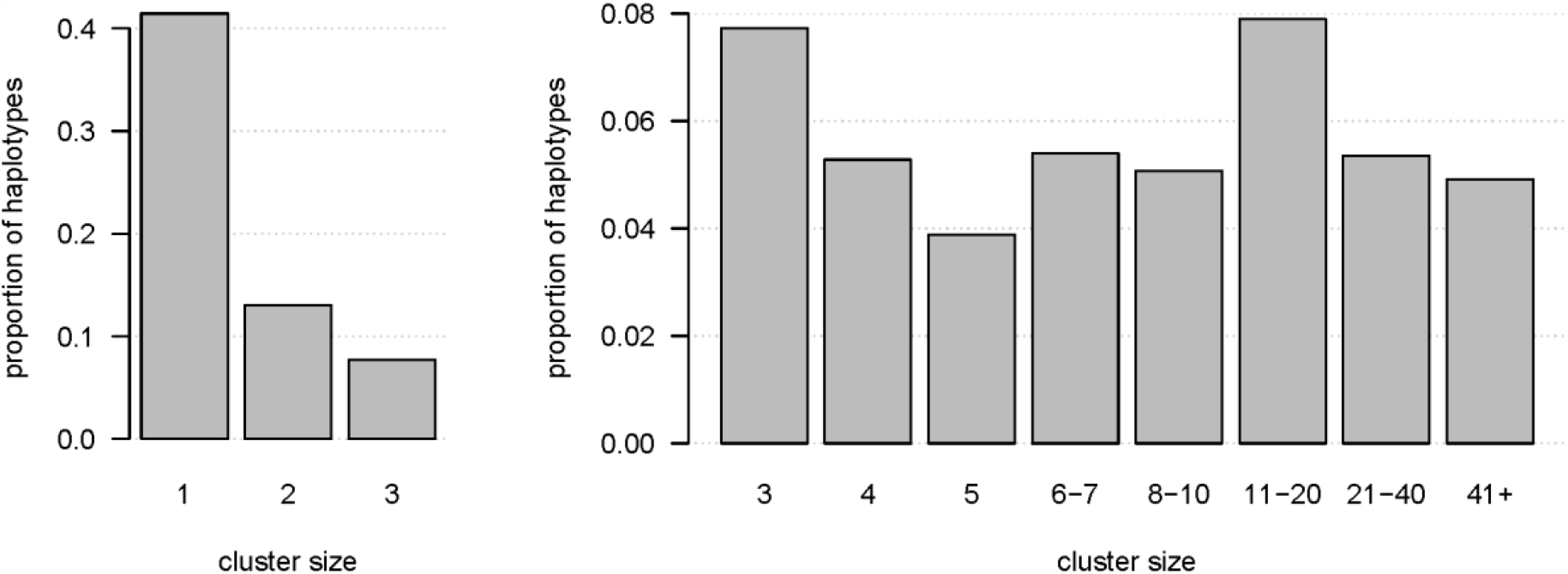
IBD cluster sizes in the UK Biobank White British autosomal sequence data. Cluster size is shown on the x-axis for cluster sizes of ≤3 in the left panel and ≥3 in the right panel. The y-axis shows the proportion of haplotypes that are in IBD clusters having that size.

We also investigated the use of smaller values of *T*. When analyzing chromosome 20 with values of *T* of 0.5 or less, we found many more spikes in the IBD rate across the chromosome compared to using *T* ≥ 0.75 (Figure S5). A spike in the IBD rate may indicate that the trim is insufficient at that location, which may be due to a short gap in the sequence data or a lower density of markers in the region. We thus recommend using a trim value of 0.75 cM or greater for human sequence data.

## Discussion

In this work, we have presented a method for inferring multi-individual IBD in biobank-scale data. The method is computationally efficient and produces simple, compact output that can be easily parsed. Analyses of simulated data show that the IBD detected with this method is highly accurate. The method scales linearly in memory use and computation time with sample size. Based on our analysis of UK Biobank and simulated data, we estimate that the method can analyze millions of genomes using ordinary computer servers.

Our approach takes a locus-centric viewpoint to IBD, rather than a segment-centric viewpoint. The combination IBD clustering and a locus-centric viewpoint enables us to output the IBD in a format that scales linearly with sample size instead of quadratically. Output of IBD clusters can be limited to positions that are separated by one or more kilobases, since IBD cluster membership does not change rapidly over these scales. Thus, the output file of IBD clusters provides a compressed storage format for IBD data.

The IBD clustering method trims the ends of pairwise IBS segments. This avoids the problem of spurious pairwise IBD perniciously interacting with IBD transitivity to create large spurious clusters of multi-individual IBD. Due to IBD segment endpoint uncertainty, the ends of detected pairwise IBD segments tend to have lower accuracy.^9^ IBS segment trimming greatly reduces false positive inference of multi-individual IBD. Our algorithm ensures that detected IBD satisfies transitivity, which is a property that is implied by the shared-ancestry definition of IBD, but that is not guaranteed by pairwise IBD detection methods.^6^

Our multi-individual IBD detection method does not currently allow for genotype errors or other causes of mismatch between IBD haplotypes. In sequence data, excluding low-frequency variants can be helpful in reducing the amount of IBD lost to genotype error.^45; 47^

We also present a method for identifying alleles that have been changed by gene conversion. Analysis of simulated data shows that the detected allele conversions have a very low false discovery rate, particularly when the trim setting for the multi-individual IBD detection is relatively high.

We applied our methods to sequence data on 125,361 individuals from the UK Biobank. We found that on average more than one-third (37.8%) of haplotypes were members of inferred IBD clusters having 4 or more haplotypes. We used these clusters to detect more than 9 million allele conversions. In contrast, the family-based deCODE study found 3237 allele conversions using a mix of SNP array data and whole genome sequence data on 7229 proband-family sets, with each proband-family set including genetic data on at least seven individuals.^36^ While more allele conversions could be found using sequence data rather than array data on all families, family-based analyses are limited in the number of meioses that can be interrogated. The average proportion of base pairs in a single meiosis that are included in gene conversion tracts is 6 × 10^−6^ and heterozygosity is less than 10^−3^ per bp in human populations. ^18; 48^ Thus, the rate of allele conversions is less than 6 × 10^−9^ per bp per meiosis, or fewer than 18 allele conversions expected per meiosis across the genome, or 36 per proband. Thus, at least 250,000 sequenced probands plus additional family members would be needed to detect the number of allele conversions detected with our method. Our IBD-based method draws from historical meioses going back tens or hundreds of generations, which enables many more allele conversions to be detected.

Our method for detecting allele conversions does not require relatives, whereas family-based analyses require multi-generational families in order to distinguish actual allele conversions from genotype error. Similarly, sperm-typing approaches have difficulty with genotype errors and are best for investigating rates of events rather than identifying individual allele conversions.^35^ Our method removes most genotype errors by requiring that the putative converted allele be carried by at least two identical-by-descent haplotypes.

The distribution of minor allele frequencies of our inferred allele conversions has the expected proportional relationship with heterozygosity. Recurrent mutations and other artifacts would not be expected to follow this relationship, indicating that most of the inferred allele conversions are real, and that the numbers of recurrent mutations and other artifacts included in the results is relatively small.

The allele conversions inferred with our method could be used to estimate the distribution of gene conversion tract lengths and the rate of gene conversion across the genome. Our method does not identify which of the two alleles in a discordant IBD cluster is the one that has been introduced through gene conversion. Since our method is based on recent meioses represented by IBD, the inferred allele conversions will have occurred within the past 100 generations or so, and thus the haplotype carrying the converted allele will have low frequency. Thus, it should be possible to determine the identity of the converted allele by considering the frequencies of the two candidate haplotypes.

## Data and Code Availability

The IBD cluster software is available from https://github.com/browning-lab/ibd-cluster. The UK Biobank data are available by application to the UK Biobank.

## Acknowledgments

This research has been conducted using the UK Biobank Resource under Application Number 19934. The methodological and analytical work performed in this study was supported by the National Human Genome Research Institute (NHGRI) under award numbers R01 HG005701 and R01 HG008359. The content is solely the responsibility of the authors and does not necessarily represent the official views of the National Institutes of Health or the UK Biobank.

## Web Resources

Beagle: https://faculty.washington.edu/browning/beagle/beagle.html

msprime: https://tskit.dev/msprime/docs/stable/intro.html

## Supplementary Methods

### Pseudocode for IBD clustering algorithm

Index the *H* haplotypes with 1, 2, …, *H*, and index the *M* markers in chromosome order with 1, 2, …, *M*. Let *W*_1_, *W*_2_, …, *W*_*J*_ be the sequence of windows. The first window begins with the first marker and ends with the first marker that is at least *L* cM away. The last marker in the windows advances by one marker per window. The first marker in a window is the closest marker that precedes that last marker by at least *L* cM. The trimmed window 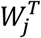 is obtained by trimming *T* cM (*T* ≤ *L*/2) from each end of *W*_*j*_.

The Positional Burrows-Wheeler Transform (PBWT) processes markers in chromosome order. At each marker, *m*, the PBWT produces two arrays of length *H*: a prefix array, *a*_*m*+1_, and a divergence array, *d*_*m*+1_.^38^

The prefix array, *a*_*m*+1_, orders the *H* haplotypes lexicographically by their reverse haplotype prefixes for the first *m* markers. The reverse haplotype prefix for the first *m* markers is the sequence of marker alleles in reverse order. Thus, if *h*[*m*] is the allele carried on haplotype *h* at the *m*-th marker, the reverse haplotype prefix for the first *m* markers is *h*[*m*], *h*[*m* − 1], …, *h*[1]. Haplotypes that have the same allele sequence in a window whose last marker is *m*, will be sorted next to each other in the PBWT prefix array *a*_*m*+1_.

For *k* > 1, the divergence array value *d*_*m*+1_[*k*] is the smallest marker, 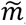, such that haplotypes *a*_*m*+1_[*k*] and *a*_*m*+1_[*k* − 1] have identical alleles between markers 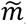 and *m* inclusive and is *m* + 1 if the haplotypes carry different alleles at marker *m*.

If marker *m* is the last marker in window *W*_*j*_, the divergence array *d*_*m*+1_ identifies pairs of adjacent haplotypes in the prefix array *a*_*m*+1_ that have the same allele sequence in the marker window. In particular, haplotypes *a*_*m*+1_[*k* − 1] and *a*_*m*+1_[*k*], have identical alleles in the window if *d*_*m*+1_[*k*] is less than or equal to the first marker in the window.

Our IBD clustering algorithm assigns all haplotypes that are confidently IBD at marker *m* to the same IBD cluster. Two haplotypes are confidently IBD at marker *m* if there is a window *W*_*j*_ in which the haplotypes have identical alleles and *m* is contained in the trimmed window 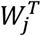. In the following pseudocode, 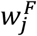 and 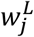 denote the first and last markers in window *W*_*j*_, and 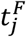 and 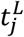 denote the first and last markers in the trimmed window 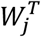.

#### IBD clustering algorithm

~~~
Inputs:
    Phased genotype data with *H* haplotypes and *M* markers.
    *L*: minimum IBS segment cM length,
    *T*: cM length trimmed from ends of IBS segment (*T* ≤ *L*/2).
// Apply PBWT to window *W*_1_, initialize IBD clusters in *W*_1_,
// and output singleton IBD clusters for markers preceding 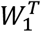.
for *m* in 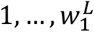:
    Compute *a*_*m*+1_ and *d*_*m*+1_
    Assign each haplotype to a singleton IBD cluster at marker *m*
    If 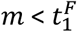:
        Output IBD clusters at marker *m*
// Merge IBD clusters in trimmed window 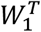
for *k* in 2, …, *H*:
    if 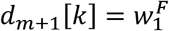:
        for each *m*′ in 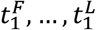:
            Merge IBD clusters at marker *m*′ containing haplotypes *a*_*m*+1_[*k* − 1] and *a*_*m*+1_[*k*]
// Process windows *W*_2_, …, *W*_*j*_. For each window:
// a) Advance the PBWT and initialize IBD clusters for last marker in the window
// b) Merge IBD clusters in the trimmed window
// c) Output IBD clusters for markers preceding the first marker in the trimmed window
for *j* in 2 … *J*:,
    let 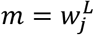
    Compute *a*_*m*+1_ and *d*_*m*+1_
    Assign each haplotype to a singleton IBD cluster at marker *m*
    for *k* in 2 … *H*:
        if 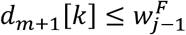:            // IBD clusters are already merged through 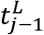
            for each *m*′ in 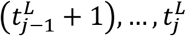:
                Merge IBD clusters at marker *m*′ containing haplotypes *a*_*m*+1_[*k* − 1] and *a*_*m*+1_[*k*]
        else if 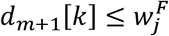:
            for each *m*′ in 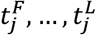:
                Merge IBD clusters at marker *m*′ containing haplotypes *a*_*m*+1_[*k* − 1] and *a*_*m*+1_[*k*]
    for *m*′ in 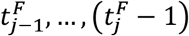:
            Output IBD clusters at marker *m*′
// Output IBD clusters for all remaining markers
for *m*′ in 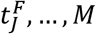:
Output IBD clusters at marker *m*′
~~~

The preceding pseudocode has low memory requirements. At any point in time, only the phased genotypes for a single marker and IBD clusters for a single marker window are stored in memory.

Our implementation of this algorithm in the ibd-cluster software package buffers the genotype data and IBD clusters in order to increase computational efficiency. This buffering does not change the scaling properties of the algorithm: memory and computation time still scale linearly with sample size.

### Determination of true IBD and allele conversions in simulated data

In order to find the true IBD segments of length at least 1 cM for a given pair of simulated haplotypes we first record the most recent common ancestor of the pair from the ancestry trees at positions spaced every 0.1 cM. If we find a sequence of at least seven adjacent interrogated positions with at most one of the positions not having the same most recent common ancestor as the others, we consider this to be an IBD segment and investigate it further to see if its length is at least 1cM. Allowing one of the positions to have a different most recent common ancestor allows for the occurrence of a gene conversion within the segment. Starting from the leftmost interrogated position with the given most recent common ancestor, we look leftward through the simulated trees at all positions until we find a position with a different most recent common ancestor, which gives the left end of the segment. We similarly look rightward from the rightmost interrogated position to find the right end of the segment. We report the IBD segment if the length is at least 1 cM. Note that if a gene conversion occurs in the first or last 0.1 cM of the segment, it will lead to a slight premature truncation of the segment which is considered acceptable for the current study.

After finding all pairwise IBD segments of length at least 1 cM, we apply transitivity to find additional smaller segments of true IBD. This application of transitivity accounts for the fact that that some true IBD segments found with our multi-individual IBD clustering will have length less than 1 cM due to transitivity (Figure 1).

Once we have found the true IBD segments as described above, we look for recent gene conversions. We do this by searching each true IBD segment for regions in which the time to most recent common ancestor differs from that in the majority of the segment. Then, at each position within each of these gene conversion regions, we check whether the two alleles carried by the two IBD haplotypes differ. If the alleles differ, we record the position as a true allele conversion.

## Supplementary Figures

**Figure S1:**
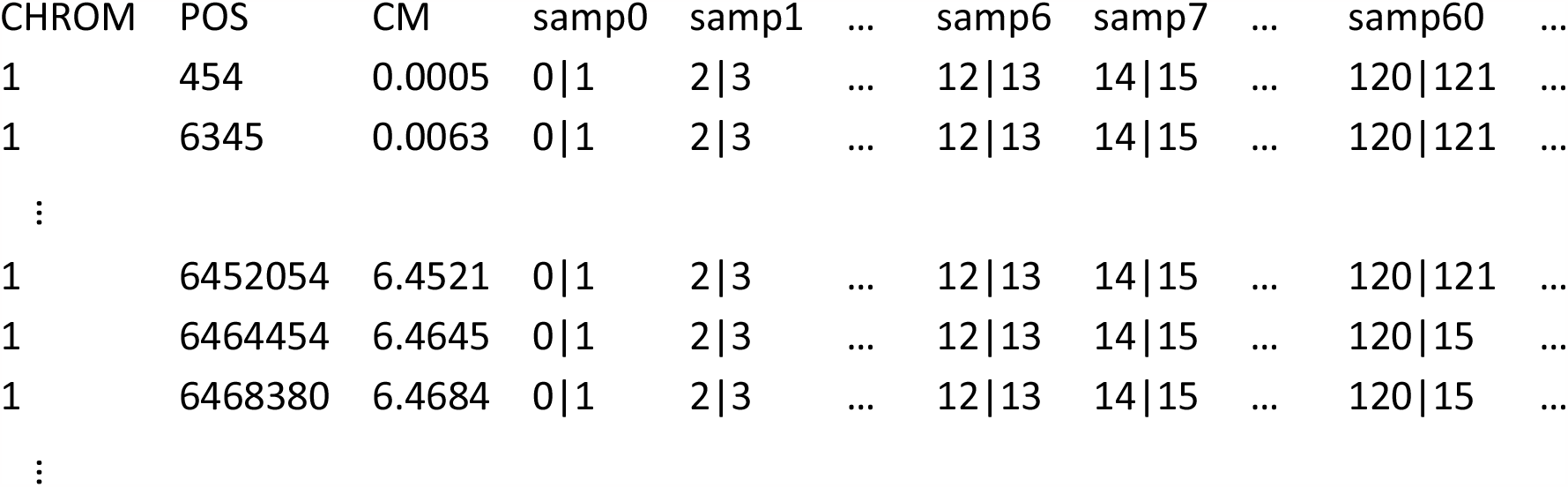
VCF-style recording of multi-individual IBD. The figure shows an extract from simulated data of the output IBD cluster file. Triple dots indicate columns/rows not shown. There is one row per position plus one header row. Fields on each row are tab delimited. There are three initial columns of genomic location information that are followed by one column per individual. Each individual at each position has two IBD cluster identifiers separated by a vertical bar. The phase of the clusters corresponds to the phase of the input genotypes. Haplotypes with the same cluster index at a position are located within the same IBD cluster at that position and are thus identical by descent. In the excerpt shown, the only identical-by-descent haplotypes are haplotype 2 of samp7 and haplotype 2 of samp60 from position 6464454 onwards. Cluster indices apply only to a single position – the same integer may correspond to different clusters at different positions.

**Figure S2:**
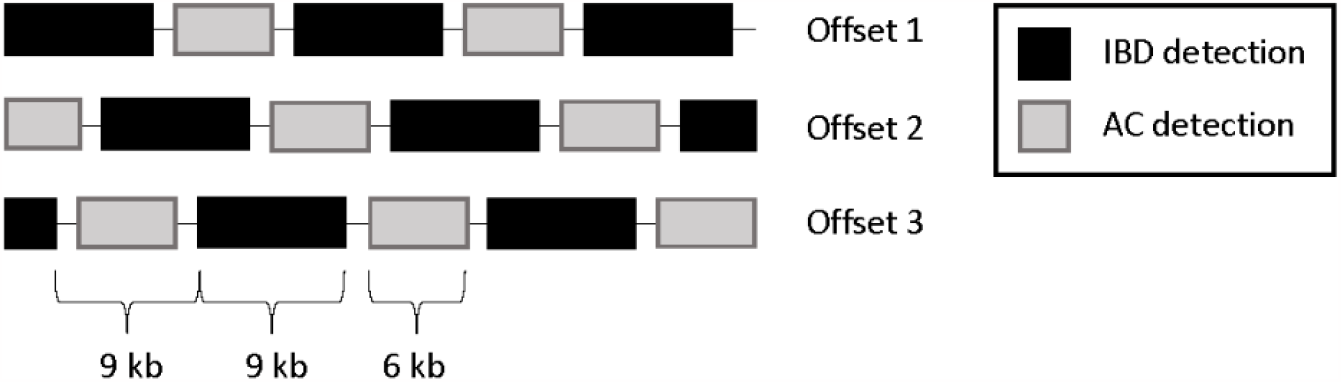
Offsets and gaps in detecting allele conversions. Three shifted sequences of IBD detection regions and allele conversion (AC) detection regions are shown in a portion of a chromosome. Since allele conversions will disrupt IBD detection, we use different sets of markers to infer IBD and to infer alleles changed by gene conversion. Each position in the chromosome is covered by one allele conversion detection region in exactly one of the three offsets.

**Figure S3:**
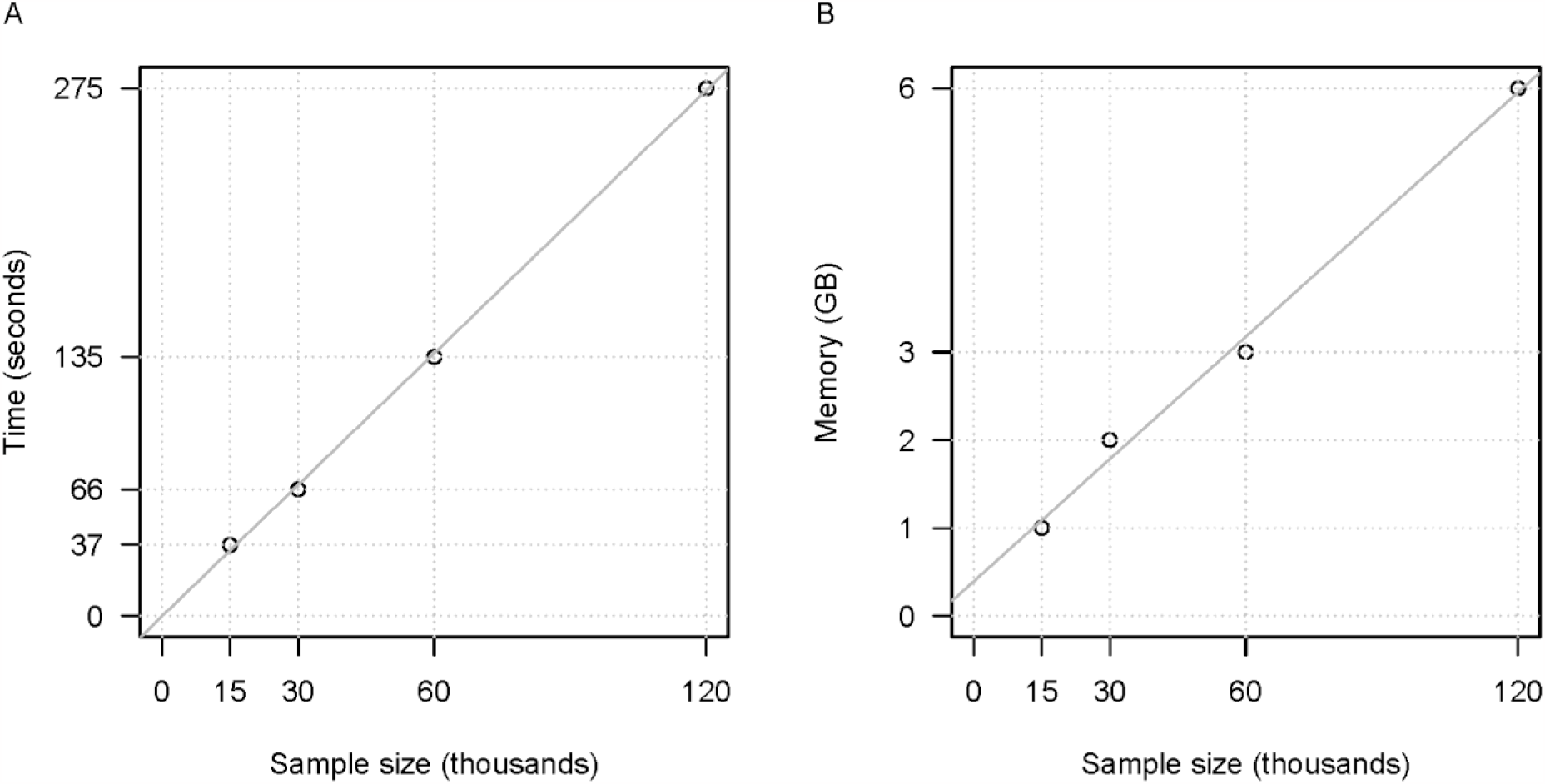
Computation time and memory requirements of the ibd-cluster software as a function of sample size for simulated data. **A**. Wall clock computation time, on an Intel Xeon Silver 4214 2.2 GHz processor with 24 CPU cores. **B**. Minimum memory needed. The Java interpreter’s maximum heap size was set using Java’s -Xmx parameter, in increments of 1 gigabase. The smallest heap size that did not result in an out-of-memory error is reported. The gray line in each panel is a least-squares regression line. The simulated data comprise one 10 Mb region as described in the main text. The analyses applied a minor allele frequency of 0.1 and a single offset of gaps as described in the main text.

**Figure S4:**
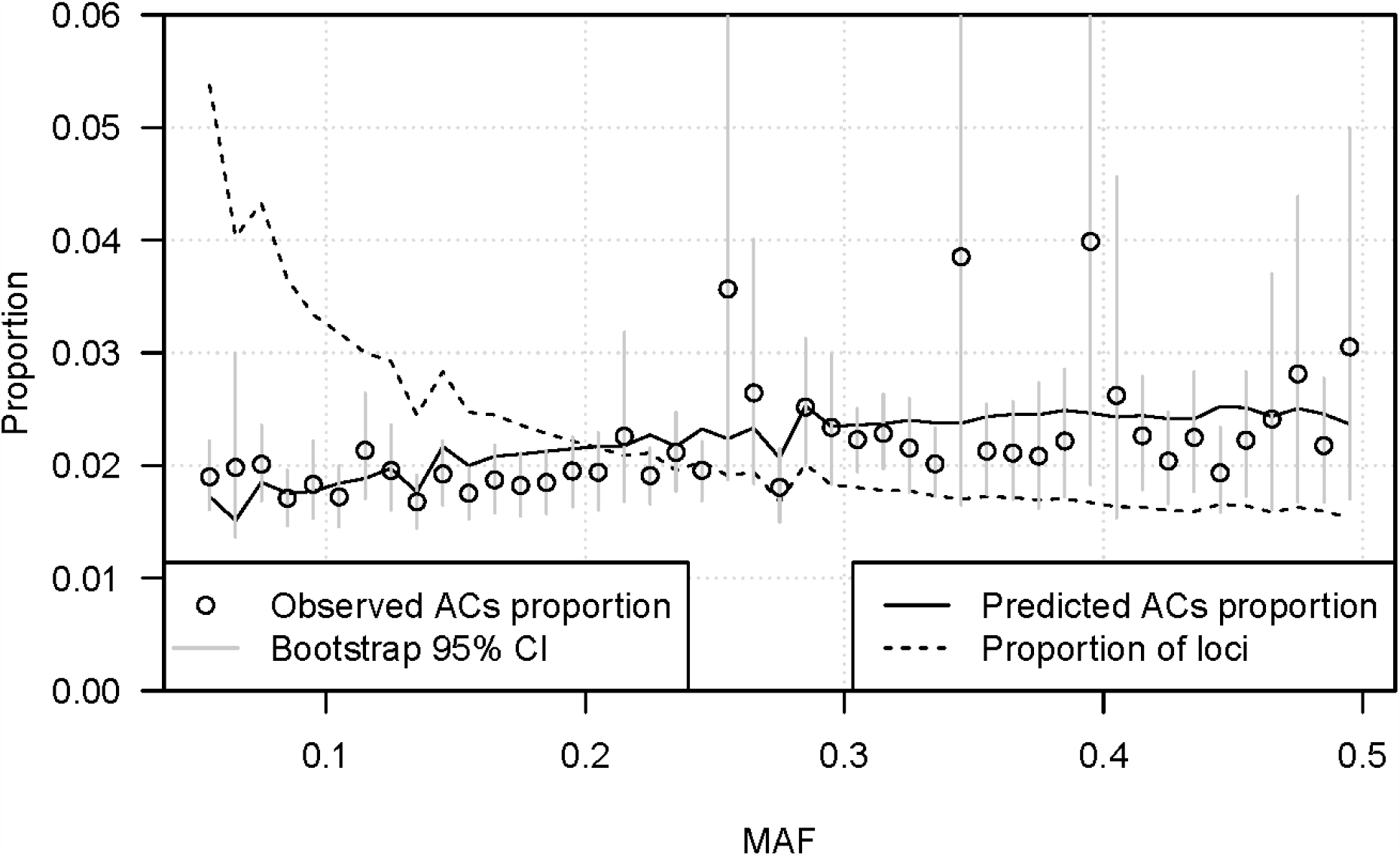
Proportion of inferred allele conversions (ACs) falling in each minor allele frequency (MAF) bin in the UK Biobank White British sequence data. MAF bins are of size 0.01, and the results are plotted at the bin midpoints. The observed proportion of inferred ACs falling within each MAF bin are shown as circles. The bootstrap 95% confidence intervals (vertical grey lines) for the AC proportions are percentile intervals from 10,000 bootstrap replicates, with bootstrapping over chromosome (chromosomes 1-22). The predicted value (solid black line) is the heterozygosity corresponding to the MAF (i.e., 2*p*(1 − *p*) where *p* is the midpoint of the MAF bin) multiplied by the number of analyzed loci with MAF within the bin, then normalized by dividing by the total of these values across all bins. The 95% confidence intervals cover the predicted value in most of the bins. The dashed line gives the proportion of analyzed loci in each MAF bin.

**Figure S5:**
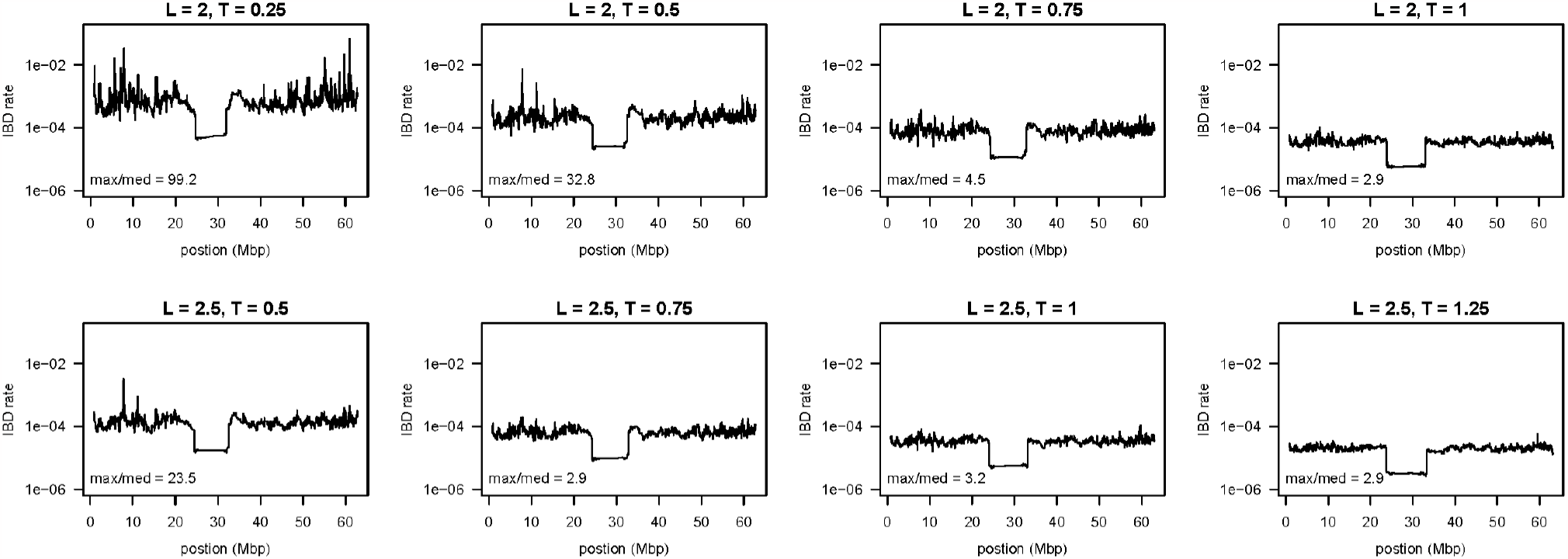
IBD rate on chromosome 20 in the UK Biobank White British sequence data with IBS length and trim values as indicated above each plot. The minimum IBS length is 2 cM in the upper row and 2.5 cM in the lower row. The maximum IBD rate divided by the median IBD rate is indicated in the lower left corner of each plot. The y-axis shows pairwise IBD rate plotted on a log scale. There is a 2.2 Mb gap in the markers at positions 26.4 Mb to 28.6 Mb due to the centromere which results in reduced IBD detection around that region.

## Supplementary Tables

**Table S1:**
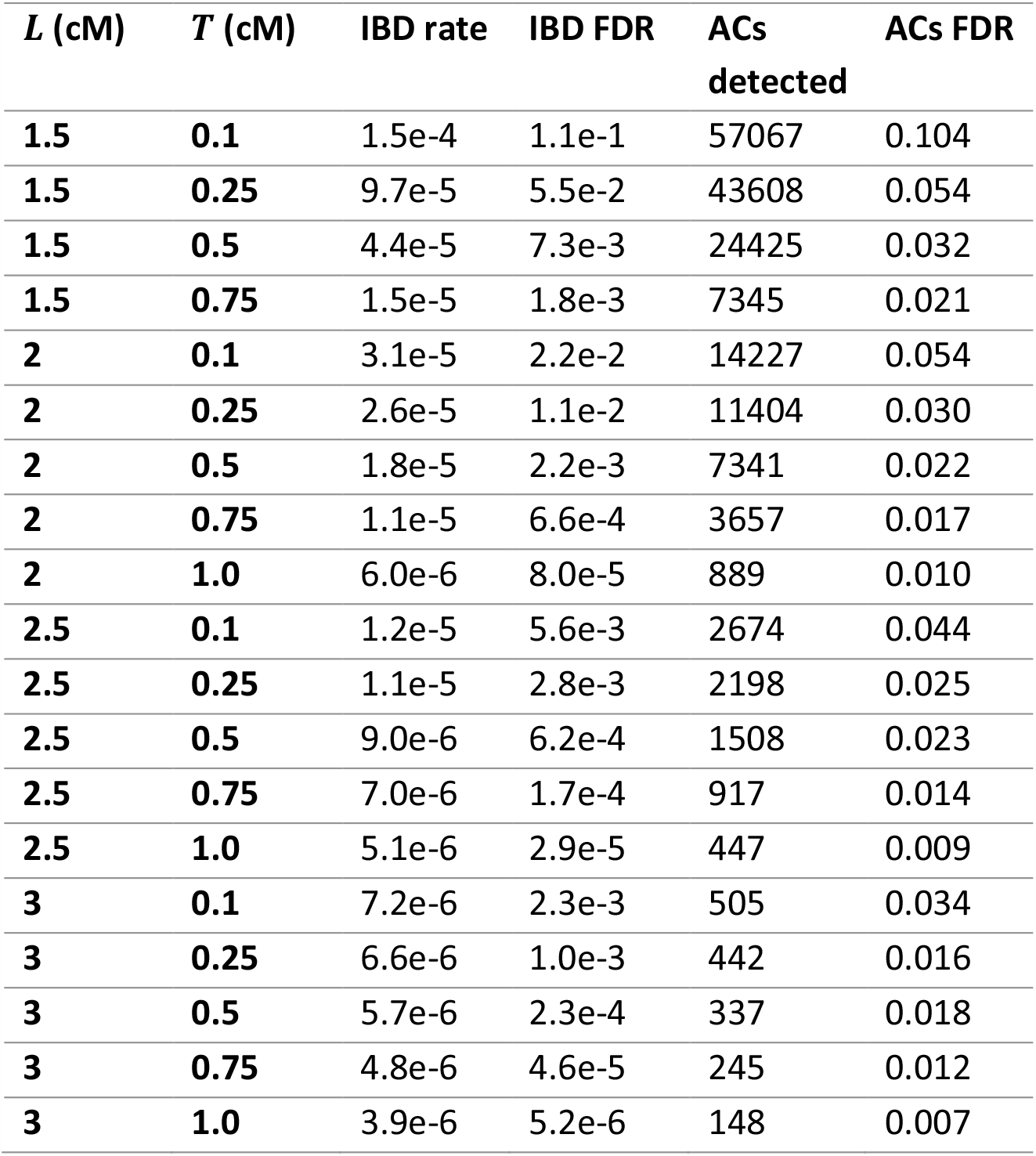
Accuracy of IBD inference and allele conversion inference in simulated data. *L* and *T* are the length and trim parameters (in cM) for the multi-individual IBD detection. The IBD rate is the proportion of pairs of haplotypes that are estimated to be IBD averaged across markers. False discovery rate (FDR) for IBD is the proportion of IBD pairs (over all markers) that are not true pairwise IBD. Numbers of estimated allele conversions (ACs) are given, and the FDR for ACs is the proportion of estimated ACs that are not true ACs. Simulated data comprise 10,000 individuals across 20 regions each of length 10 Mb.

| $L$ (cM) | $T$ (cM) | IBD rate | IBD FDR | ACs<br>detected | ACs FDR |
| --- | --- | --- | --- | --- | --- |
| 1.5 | 0.1 | 1.5e-4 | 1.1e-1 | 57067 | 0.104 |
| 1.5 | 0.25 | 9.7e-5 | 5.5e-2 | 43608 | 0.054 |
| 1.5 | 0.5 | 4.4e-5 | 7.3e-3 | 24425 | 0.032 |
| 1.5 | 0.75 | 1.5e-5 | 1.8e-3 | 7345 | 0.021 |
| 2 | 0.1 | 3.1e-5 | 2.2e-2 | 14227 | 0.054 |
| 2 | 0.25 | 2.6e-5 | 1.1e-2 | 11404 | 0.030 |
| 2 | 0.5 | 1.8e-5 | 2.2e-3 | 7341 | 0.022 |
| 2 | 0.75 | 1.1e-5 | 6.6e-4 | 3657 | 0.017 |
| 2 | 1.0 | 6.0e-6 | 8.0e-5 | 889 | 0.010 |
| 2.5 | 0.1 | 1.2e-5 | 5.6e-3 | 2674 | 0.044 |
| 2.5 | 0.25 | 1.1e-5 | 2.8e-3 | 2198 | 0.025 |
| 2.5 | 0.5 | 9.0e-6 | 6.2e-4 | 1508 | 0.023 |
| 2.5 | 0.75 | 7.0e-6 | 1.7e-4 | 917 | 0.014 |
| 2.5 | 1.0 | 5.1e-6 | 2.9e-5 | 447 | 0.009 |
| 3 | 0.1 | 7.2e-6 | 2.3e-3 | 505 | 0.034 |
| 3 | 0.25 | 6.6e-6 | 1.0e-3 | 442 | 0.016 |
| 3 | 0.5 | 5.7e-6 | 2.3e-4 | 337 | 0.018 |
| 3 | 0.75 | 4.8e-6 | 4.6e-5 | 245 | 0.012 |
| 3 | 1.0 | 3.9e-6 | 5.2e-6 | 148 | 0.007 |

## Literature cited

1. Gusev, A., Lowe, J.K., Stoffel, M., Daly, M.J., Altshuler, D., Breslow, J.L., Friedman, J.M., and Pe’er, I. (2009). Whole population, genome-wide mapping of hidden relatedness. Genome Res 19, 318–326.

2. Browning, S.R., and Browning, B.L. (2012). Identity by descent between distant relatives: detection and applications. Annual Review of Genetics 46, 617–633.

3. Sticca, E.L., Belbin, G.M., and Gignoux, C.R. (2021). Current developments in detection of identity-by-descent methods and applications. Frontiers in Genetics 12, 722602.

4. Te Meerman, G.J., Van Der Meulen, M.A., and Sandkuijl, L.A. (1995). Perspectives of identity by descent (IBD) mapping in founder populations. Clinical & Experimental Allergy 25, 97–102.

5. Purcell, S., Neale, B., Todd-Brown, K., Thomas, L., Ferreira, M.A.R., Bender, D., Maller, J., Sklar, P., de Bakker, P.I.W., Daly, M.J., et al. (2007). PLINK: A tool set for whole-genome association and population-based linkage analyses. American Journal of Human Genetics 81, 559–575.

6. Gusev, A., Kenny, E.E., Lowe, J.K., Salit, J., Saxena, R., Kathiresan, S., Altshuler, D.M., Friedman, J.M., Breslow, J.L., and Pe’er, I. (2011). DASH: a method for identical-by-descent haplotype mapping uncovers association with recent variation. American Journal of Human Genetics 88, 706–717.

7. Browning, S.R., and Thompson, E.A. (2012). Detecting rare variant associations by identity-by-descent mapping in case-control studies. Genetics 190, 1521–1531.

8. Albrechtsen, A., Moltke, I., and Nielsen, R. (2010). Natural selection and the distribution of identity-by-descent in the human genome. Genetics 186, 295–308.

9. Browning, S.R., and Browning, B.L. (2020). Probabilistic Estimation of Identity by Descent Segment Endpoints and Detection of Recent Selection. American Journal of Human Genetics 107, 895–910.

10. Huff, C.D., Witherspoon, D.J., Simonson, T.S., Xing, J., Watkins, W.S., Zhang, Y., Tuohy, T.M., Neklason, D.W., Burt, R.W., Guthery, S.L., et al. (2011). Maximum-likelihood estimation of recent shared ancestry (ERSA). Genome Research 21, 768–774.

11. Henn, B.M., Hon, L., Macpherson, J.M., Eriksson, N., Saxonov, S., Pe’er, I., and Mountain, J.L. (2012). Cryptic distant relatives are common in both isolated and cosmopolitan genetic samples. PLoS One 7, e34267.

12. Seidman, D.N., Shenoy, S.A., Kim, M., Babu, R., Woods, I.G., Dyer, T.D., Lehman, D.M., Curran, J.E., Duggirala, R., Blangero, J., et al. (2020). Rapid, Phase-free Detection of Long Identity-by-Descent Segments Enables Effective Relationship Classification. American Journal of Human Genetics 106, 453–466.

13. Zhou, Y., Browning, S.R., and Browning, B.L. (2020). IBDkin: fast estimation of kinship coefficients from identity by descent segments. Bioinformatics 36, 4519–4520.

14. Ralph, P., and Coop, G. (2013). The geography of recent genetic ancestry across Europe. PLoS Biol 11, e1001555.

15. Palamara, P.F., Lencz, T., Darvasi, A., and Pe’er, I. (2012). Length distributions of identity by descent reveal fine-scale demographic history. American Journal of Human Genetics 91, 809–822.

16. Browning, S.R., and Browning, B.L. (2015). Accurate non-parametric estimation of recent effective population size from segments of identity by descent. Am J Hum Genet 97, 404–418.

17. Palamara, P.F., and Pe’er, I. (2013). Inference of historical migration rates via haplotype sharing. Bioinformatics 29, i180–188.

18. Palamara, P.F., Francioli, L.C., Wilton, P.R., Genovese, G., Gusev, A., Finucane, H.K., Sankararaman, S., Genome of the Netherlands Consortium, Sunyaev, S.R., de Bakker, P.I.W., et al. (2015). Leveraging Distant Relatedness to Quantify Human Mutation and Gene-Conversion Rates. American Journal of Human Genetics 97, 775–789.

19. Tian, X., Browning, B.L., and Browning, S.R. (2019). Estimating the Genome-wide Mutation Rate with Three-Way Identity by Descent. Am J Hum Genet 105, 883–893.

20. Zhou, Y., Browning, B.L., and Browning, S.R. (2020). Population-Specific Recombination Maps from Segments of Identity by Descent. American Journal of Human Genetics 107, 137–148.

21. Naseri, A., Yue, W., Zhang, S., and Zhi, D. (2023). Fast inference of genetic recombination rates in biobank scale data. Genome Research, gr. 277676.277123.

22. Browning, S.R. (2008). Estimation of pairwise identity by descent from dense genetic marker data in a population sample of haplotypes. Genetics 178, 2123–2132.

23. Kong, A., Masson, G., Frigge, M.L., Gylfason, A., Zusmanovich, P., Thorleifsson, G., Olason, P.I., Ingason, A., Steinberg, S., Rafnar, T., et al. (2008). Detection of sharing by descent, long-range phasing and haplotype imputation. Nature Genetics 40, 1068–1075.

24. Browning, S.R., and Browning, B.L. (2010). High-resolution detection of identity by descent in unrelated individuals. Am J Hum Genet 86, 526–539.

25. Abney, M., and Han, L. (2011). Identity by Descent Estimation With Dense Genome-Wide Genotype Data. Genetic Epidemiology 35, 557–567.

26. Dimitromanolakis, A., Paterson, A.D., and Sun, L. (2019). Fast and accurate shared segment detection and relatedness estimation in un-phased genetic data via TRUFFLE. The American Journal of Human Genetics 105, 78–88.

27. Naseri, A., Liu, X., Tang, K., Zhang, S., and Zhi, D. (2019). RaPID: ultra-fast, powerful, and accurate detection of segments identical by descent (IBD) in biobank-scale cohorts. Genome Biol 20, 1–15.

28. Shemirani, R., Belbin, G.M., Avery, C.L., Kenny, E.E., Gignoux, C.R., and Ambite, J.L. (2021). Rapid detection of identity-by-descent tracts for mega-scale datasets. Nat Commun 12, 3546.

29. Tian, X., Cai, R., and Browning, S.R. (2022). Estimating the genome-wide mutation rate from thousands of unrelated individuals. The American Journal of Human Genetics 109, 2178–2184.

30. Qiao, Y., Sannerud, J., Basu-Roy, S., Hayward, C., and Williams, A.L. (2019). Distinguishing pedigree relationships using multi-way identical by descent sharing and sex-specific genetic maps. BioRxiv, 753343.

31. Danecek, P., Auton, A., Abecasis, G., Albers, C.A., Banks, E., DePristo, M.A., Handsaker, R.E., Lunter, G., Marth, G.T., Sherry, S.T., et al. (2011). The variant call format and VCFtools. Bioinformatics 27, 2156–2158.

32. Qian, Y., Browning, B.L., and Browning, S.R. (2014). Efficient clustering of identity-by-descent between multiple individuals. Bioinformatics 30, 915–922.

33. Shemirani, R., Belbin, G.M., Burghardt, K., Lerman, K., Avery, C.L., Kenny, E.E., Gignoux, C.R., and Ambite, J.L. (2023). Selecting Clustering Algorithms for Identity-By-Descent Mapping. Pac Symp Biocomput 28, 121–132.

34. Williams, A.L., Genovese, G., Dyer, T., Altemose, N., Truax, K., Jun, G., Patterson, N., Myers, S.R., Curran, J.E., Duggirala, R., et al. (2015). Non-crossover gene conversions show strong GC bias and unexpected clustering in humans. Elife 4.

35. Jeffreys, A.J., and May, C.A. (2004). Intense and highly localized gene conversion activity in human meiotic crossover hot spots. Nature genetics 36, 151–156.

36. Halldorsson, B.V., Hardarson, M.T., Kehr, B., Styrkarsdottir, U., Gylfason, A., Thorleifsson, G., Zink, F., Jonasdottir, A., Jonasdottir, A., Sulem, P., et al. (2016). The rate of meiotic gene conversion varies by sex and age. Nature Genetics 48, 1377–1384.

37. Gay, J., Myers, S., and McVean, G. (2007). Estimating meiotic gene conversion rates from population genetic data. Genetics 177, 881–894.

38. Durbin, R. (2014). Efficient haplotype matching and storage using the positional Burrows-Wheeler transform (PBWT). Bioinformatics 30, 1266–1272.

39. Cormen, T.H., Leiserson, C.E., Rivest, R.L., and Stein, C. (2009). Introduction to algorithms.(MIT press).

40. Baumdicker, F., Bisschop, G., Goldstein, D., Gower, G., Ragsdale, A.P., Tsambos, G., Zhu, S., Eldon, B., Ellerman, E.C., Galloway, J.G., et al. (2022). Efficient ancestry and mutation simulation with msprime 1.0. Genetics 220.

41. Taliun, D., Harris, D.N., Kessler, M.D., Carlson, J., Szpiech, Z.A., Torres, R., Taliun, S.A.G., Corvelo, A., Gogarten, S.M., Kang, H.M., et al. (2021). Sequencing of 53,831 diverse genomes from the NHLBI TOPMed Program. Nature 590.

42. Browning, B.L., Tian, X., Zhou, Y., and Browning, S.R. (2021). Fast two-stage phasing of large-scale sequence data. The American Journal of Human Genetics 108, 1880–1890.

43. Bycroft, C., Freeman, C., Petkova, D., Band, G., Elliott, L.T., Sharp, K., Motyer, A., Vukcevic, D., Delaneau, O., and O’Connell, J. (2018). The UK Biobank resource with deep phenotyping and genomic data. Nature 562, 203–209.

44. Halldorsson, B.V., Eggertsson, H.P., Moore, K.H.S., Hauswedell, H., Eiriksson, O., Ulfarsson, M.O., Palsson, G., Hardarson, M.T., Oddsson, A., Jensson, B.O., et al. (2022). The sequences of 150,119 genomes in the UK Biobank. Nature 607, 732–740.

45. Browning, B.L., and Browning, S.R. (2023). Statistical phasing of 150,119 sequenced genomes in the UK Biobank. The American Journal of Human Genetics 110, 161–165.

46. Halldorsson, B.V., Palsson, G., Stefansson, O.A., Jonsson, H., Hardarson, M.T., Eggertsson, H.P., Gunnarsson, B., Oddsson, A., Halldorsson, G.H., Zink, F., et al. (2019). Characterizing mutagenic effects of recombination through a sequence-level genetic map. Science 363.

47. Zhou, Y., Browning, S.R., and Browning, B.L. (2020). A Fast and Simple Method for Detecting Identity-by-Descent Segments in Large-Scale Data. American Journal of Human Genetics 106, 426–437.

48. Mallick, S., Li, H., Lipson, M., Mathieson, I., Gymrek, M., Racimo, F., Zhao, M., Chennagiri, N., Nordenfelt, S., Tandon, A., et al. (2016). The Simons Genome Diversity Project: 300 genomes from 142 diverse populations. Nature 538, 201–206.

